# Eigenvector-based community detection for identifying information hubs in neuronal networks

**DOI:** 10.1101/457143

**Authors:** Ruaridh Clark, Malcolm Macdonald

## Abstract

Eigenvectors of networked systems are known to reveal central, well-connected, network vertices. Here we expand upon the known applications of eigenvectors to define well-connected communities where each is associated with a prominent vertex. This form of community detection provides an analytical approach for analysing the dynamics of information flow in a network. When applied to the neuronal network of the nematode *Caenorhabditis elegans*, known circuitry can be identified as separate eigenvector-based communities. For the macaque’s neuronal network, community detection can expose the hippocampus as an information hub; this result contradicts current thinking that the analysis of static graphs cannot reveal such insights. The application of community detection on a large scale human connectome (~1.8 million vertices) reveals the most prominent information carrying pathways present during a magnetic resonance imaging scan. We demonstrate that these pathways can act as an effective unique identifier for a subject’s brain by assessing the number of matching pathways present in any two connectomes.

**Author summary:** The dynamic response of a network to stimulus can be understood by investigating that system’s eigenvectors. The eigenvectors highlight the most prominent nodes; those that are either a major source or destination for information in the network. Moreover by defining a coordinate system based on multiple eigenvectors, the most prominent communities can be detected with the most prominent node detected alongside those in the community that funnel information towards it. These methods are applied to a variety of brain networks to highlight the circuitry present in a flatworm *(Caenorhabditis elegans)*, the macaque and human subjects. Static graphs representing the connectomes are analysed to provide insights that were previously believed to only be detectable by numerically modelling information flow. Finally, we discovered that brain networks created for human subjects at different times can be identified as belonging to the same subject by investigating the similarity of the prominent communities.

## Introduction

Understanding the brain’s function is a major pursuit of humanity, with mapping and comprehending the human connectome the final goal. The work contained herein develops tools that facilitate comprehension of the vast and, on the surface, incomprehensible network of the brain with, in the case of humans, an estimated 100 billion neurons and 100 trillion synaptic connections [1]. Our ability to map neurons and their connections is limited but ever improving. The magnetic resonance imaging (MRI) scans considered herein achieve a 1 mm resolution for *in vivo* human subjects [2] but 0.1 mm has been achieved with *in vivo* diffusion MRI data [3]. Improving scan resolutions will enable higher fidelity connectomes and in turn require analytical tools that can cope with the increase in scale. These detailed scans can be converted into graphs through pipelines that produce consistent connectomes allowing studies and comparisons [4]. Connectome analysis has often been constrained to small graphs (< 1000 vertices) with the results of these studies compared with existing intuitions and knowledge gained through experimentation [5], [6]. These previous studies on small connectomes have identified influential regions by performing numerical flow simulations of information travelling throughout the brain, but such an approach would likely be intractable on large graphs containing millions of nodes.

The application of graph theory on a networked system produces an abstract representation, i.e. a graph comprising of edges and vertices, that can be analysed to better understand the movement of information. It has been argued that graph theory has some blindspots when considering the dynamics of information flow, see [6], with centrality measures unable to achieve the same insights as numerical methods. Eigenvector centrality, see [7], is a metric that captures important information on a graph’s dynamics, in particular it highlights the vertices through which information most frequently passes with this capability famously underpinning Google’s PageRank development [8].

Spectral analysis, and eigenvectors in particular, are a cornerstone of the developments herein with eigenvector centrality adapted and expanded upon. This expansion is based on incorporating multiple eigenvectors to produce a dynamics-based method for community detection. A common approach for community detection is to focus on the static topology of the graph, i.e. the distribution of edges in a graph. One such example is Leicht-Newman community detection for directed graph [9] that compares the density of edges with that of a graph where edges are distributed randomly to determine whether clustering and, hence, community division is present.

In contrast to Leicht-Newman’s approach, dynamics-based community detection can reveal communities of information flow that form in the presence of stimulus. These communities are likely to be influenced by clustering but they are also affected by the source and destination of information in the graph. Simulations of information flow are an accepted approach for uncovering these dynamic processes [6, 10]. But analytical solutions may be required if connectomes grow to capture the activity of billions of neurons.

## Results

### C. Elegans Connectome

The *Caenorhabditis elegans* is a non-parasitic nematode that is transparent and unsegmented with a long cylindrical body shape that grows to about 1 mm in length. A wiring diagram of the *C. elegans* nervous system was updated by Varshney et al. to contain 279 somatic neurons [5]. The full wiring diagram includes chemical synapses, gap junctions, and neuromuscular junctions. Varshney et al. examined the undirected electrical gap junction network (containing 890 edges with a giant component of 248 vertices) and analysed the network’s eigenvectors to identify *circuits* that were highlighted in previous experimental studies. One such example was the identification of two distinct *circuits* by the eigenvector, *v*_*L*3,_ associated with *λ*_3_ where *v_L_* indicates an eigenvector of the Laplacian matrix. The *circuit* designation of vertices being determined by the sign of their entry in *v*_*L*3._ Here we extend these intuitions from a single dimension to multiple by examining the eigenvector entries *v*_*L*2,_ *v*_*L*3_ and *v*_*L*4_ as displayed in Fig. 1. This approach is the basis of the community detection method Communities of Dynamic Response (CDR) detailed in Algorithm 1. The *circuits* or, as they will be referred to from hereon, communities identified by Varshney et al. [5] are clearly visible in Fig. 1 and labelled as members of the positive *v_L_*_3_ community (orange) or the negative *v_L_*_3_ community (yellow).

**Fig 1.**
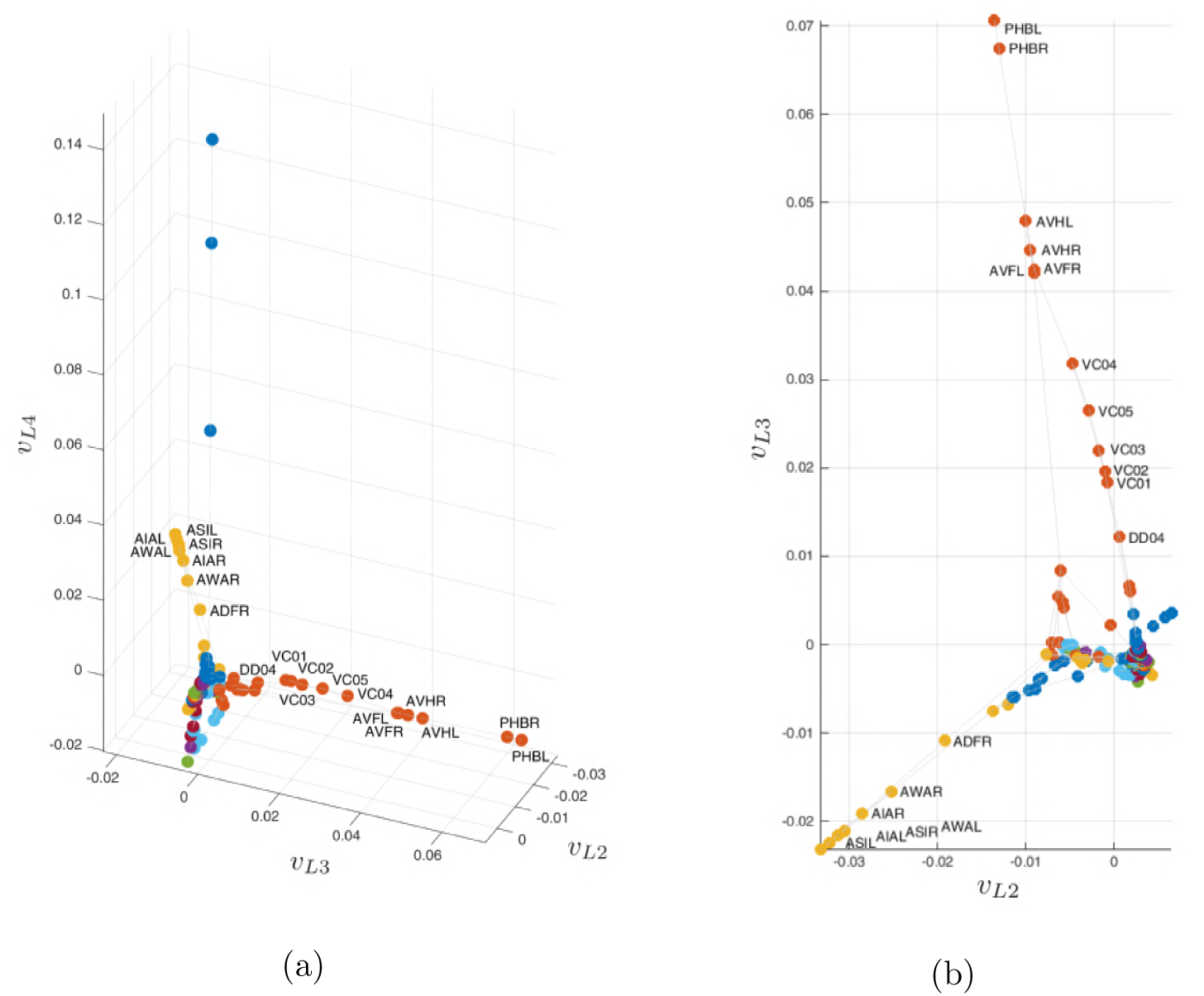
Visualisation of vertex placement in eigenvector space, where *v*_*L*2_, *v*_*L*3_ and *v*_*L*4_ are associated with *λ*_2_, *λ*_3_ and *λ*_4_ of the Laplacian matrix for the undirected electrical junction network of the *C. elegans*. Community designation (see Algorithm 1) is noted using vertex colour.

The one dimensional approach of Varshney et al. is only effective in certain cases as it can only ever identify two communities. For example, *v_L_*_6_ presented in Fig. 2 (b) could be employed to separate the vertices into two groups, but by employing CDR with three dimensions (*v_L_*_5_, *v_L_*_6_ and *v_L_*_7),_ shown in Fig. 2 (a), a more nuanced picture emerges where multiple communities are present. The community found by [5] from *v*_*L*3_ is marked with yellow vertices in Fig 1 and contains ASIL, AIAL, ASIR, AWAL, AIAR, AWAR & ADFR. This same community is not identifiable using *v_L_*_6_ in isolation but is depicted in light blue vertices in Fig. 2, which supports the validity of the community designations detailed by the CDR method.

**Fig 2.**
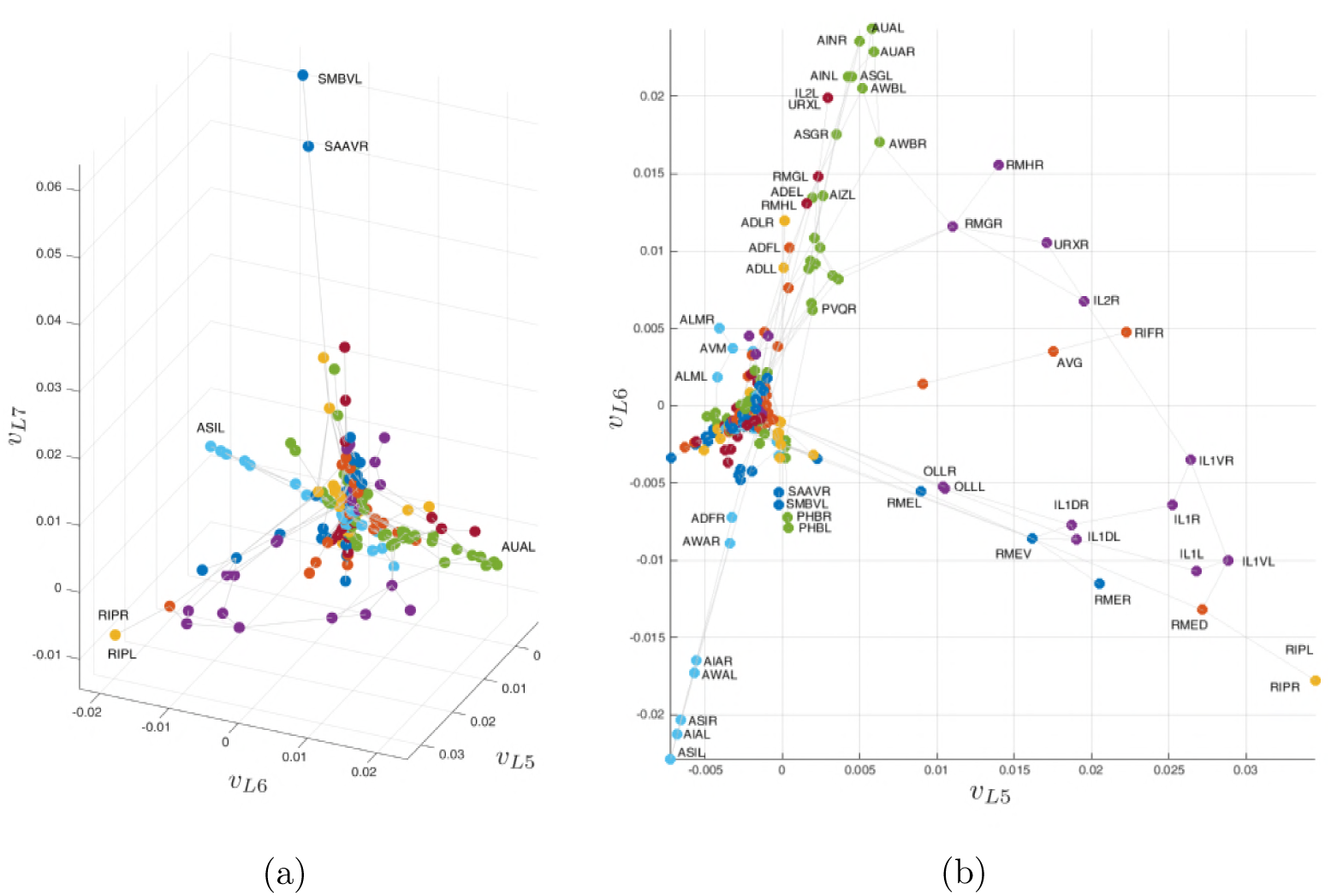
Visualisation of vertex placement in eigenvector space, where *v*_*L*5_, *v*_*L*6_ and *v*_*L*7_ are associated with *λ*_5_, *λ*_6_ and *λ*_7_ of the Laplacian matrix for the undirected electrical junction network of the C. elegans. Community designation (see Algorithm 1) is noted using vertex colour.

### Macaque Connectome

The CDR relies on eigenvectors to detect communities in a network. These eigenvectors capture the dynamics of the system and, hence, the CDR can also be exploited to determine prominent vertices in a network.

Mišić et al. [6] state that the hippocampus (CA1) of the macaque has long been known to neuroscientists to hold an influential role in the brain’s decision making architecture. Mišić et al. cite a number of graph theory based studies that have failed to identify the hippocampus as an important hub for humans [11–13] and for the macaque [14–17]. Stating that these *“analyses of anatomical and functional whole-brain networks have largely failed to demonstrate the topological centrality of the hippocampus.”* The conclusion of that study was *“the functional capacity of a given region or subnetwork cannot be fully discerned by only analyzing the static structural connectivity of the brain”*. Functional capacity is understood to be an assessment of how effectively a region can carry out a given function. In the case of the hippocampus, this would be an assessment of its ability to receive information from across the whole network.

The CoCoMac database [18] supplied the connectome used by Mišić et al. [6], which contained 242 vertices representing neuronal areas with each edge carrying equal weighting. The CDR will demonstrate that, contrary to the claims of [6], the influence of the hippocampus (CA1) can be discerned by analysing the structural connectivity of the brain.

TFM is most prominent vertex in the macaque connectome, as shown in Fig. 3, and it belongs to a community of two vertices (TFM and CA1) that are highlighted in pink. CA1 is therefore part of the most prominent pathway for information in this connectome. CA1’s influence is further enhanced by being connected to TFL, which is the second most prominent vertex according to *v_L_*_1._ CA1 receives one of the two outgoing connections from TFL, whilst also being the only receptor of information from TFM. The reason why TFM and TFL have large values of *v*_*L*1_ is that they are bottlenecks for information in the network with information from across the graph arriving at these two nodes and only having three paths to choose from, two of which lead to the CA1. A large indegree would be an intuitive identifier of prominent vertices, but TFL has only the 20^th^ highest indegree in the network. The outdegree is also a critical factor, as a vertex with a high outdegree and indegree will pass on much of the information it receives and not act as a bottleneck. This claim is supported by the ratio of outdegree to indegree (*O* : *I*) being significantly higher for TFM and TFL than any other vertex, where the ratio is 34 and 20.5 respectively. The next closest vertices, when sorted by *O* : *I* ratio, are as follows: DG (ratio of 9), 28m (8), D9 (7.7), TSA (7.3), and M2-HL (6). These vertices are clearly prominent in Fig. 3 but the order according to the *O* : *I* ratio differs from *v*_*L*1_.

**Fig 3.**
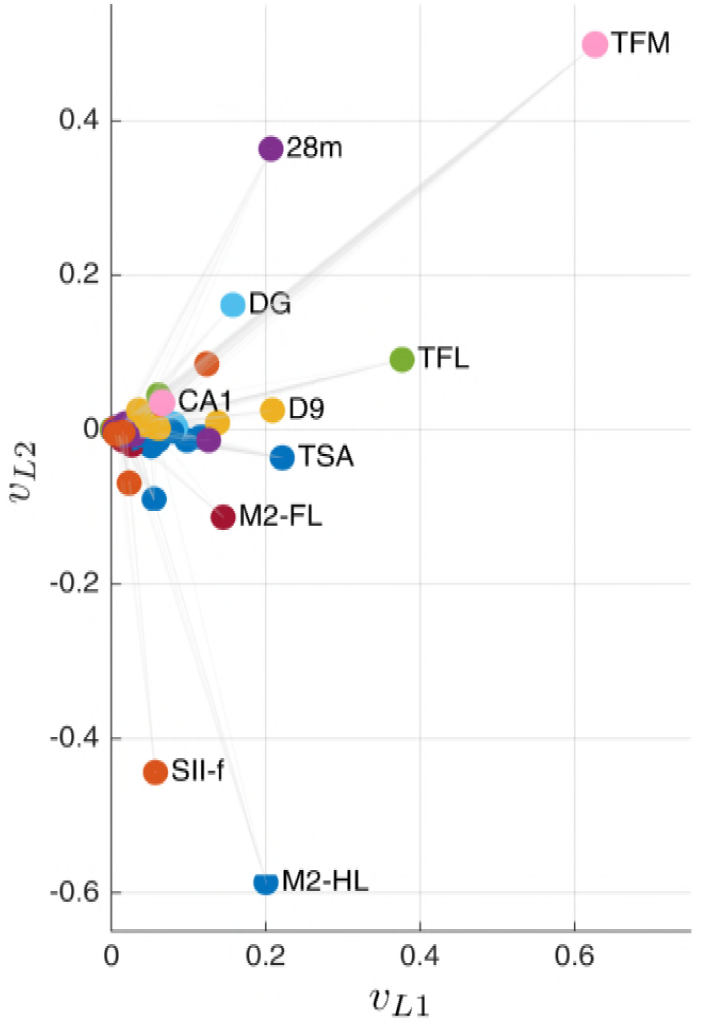
*v*_*L*1_ and *v*_*L*2_ are the first and second left eigenvectors associated with *λ*_1_ and *λ*_2_ of the Laplacian matrix for the CoCoMac network. Community designation according to Algorithm 1 is noted using vertex colour.

The CoCoMac network employs a uniform weighting for all edges. A macaque’s brain will have variable weights for the edges between different neuronal areas. Therefore, it is possible that if edge weights were known for this network, and the edge between TFM and CA1 had a large weighting, then the hippocampus (CA1) could be the largest element of *v_L_*_1._ But given the lack of edge weighting information present, an intuition is applied for this network that the highest *v*_*L*1_ vertices are the main information collators (bottlenecks). These collator vertices tend to have relatively low outdegree and pass information onto an influential region. This intuition can be tested by defining the collation vertices as prominent vertices (PV) and vertices they pass information on to as Outgoing Connection vertices (OCN). Table 1 details the PVs and OCNs for a number of prominent vertices from Fig 3. A clear example of an OCN as a known influential region is the M2-HL vertex as the PV that is connected to the M1-HL as the OCN. In this case, the supplementary motor cortex (M2-HL) is acting as the information collator that then provides information to the primary motor cortex (M1-HL), which is the most influential region for motor control.

**Table 1.** Prominent vertices from Fig 3.

| Prominent vertex (PV) |  | Outgoing Connection vertex (OCN) |  |
| --- | --- | --- | --- |
| Acronym | Merged Brain Region | Acronym | Merged Brain Region |
| TFM | Temporal area TF (medial part) | CA1 | Hippocampus |
| TFL | Temporal area TF (lateral part) | CA1, Pros. | Hippocampus, Prosubiculum |
| TSA | Transitional sensory area | 23c, 31, PECg | Area 23c, Area 31, Parietal area PE (cingulate part) |
| D9 | Dorsal area 9 | 32, 14, M9 | Area 32, Orbitofrontal area 14, Medial area 9 |
| 28m | Medial entorhinal cortex | TG | Temporopolar area TG |
| M2-HL | Supplementary motor cortex M2, hindlimb area | M1-HL | Primary motor cortex M1, hindlimb area |
| DG | Dentate gyrus | ENT | Entorhinal cortex |

A numerical flow model applied to the network in [6] produced a list of the most traversed edges in the graph, which are detailed in Table 2 alongside the *v*_*L*1_ ranking of the vertices at either end of the edge. It is evident from the table that the outgoing connection vertex is ranked highly, and for the most part in *v*_*L*1_ order. The only change in *v*_*L*1_ order, for the outgoing vertices, could be attributed to vertex D9 having two highly traversed edges. Whilst the lower ranked neuronal area 28m only has the one and so more traffic accumulates on that edge. This lends further support to the claim that the vertices ranked by *v*_*L*1_ can be viewed as bottlenecks for information from across the whole network, as many of the PV vertices funnel information in bulk to certain locations. Again it is worth noting that insights into the role of PV vertices are limited by the absence of edge weight information.

**Table 2.**
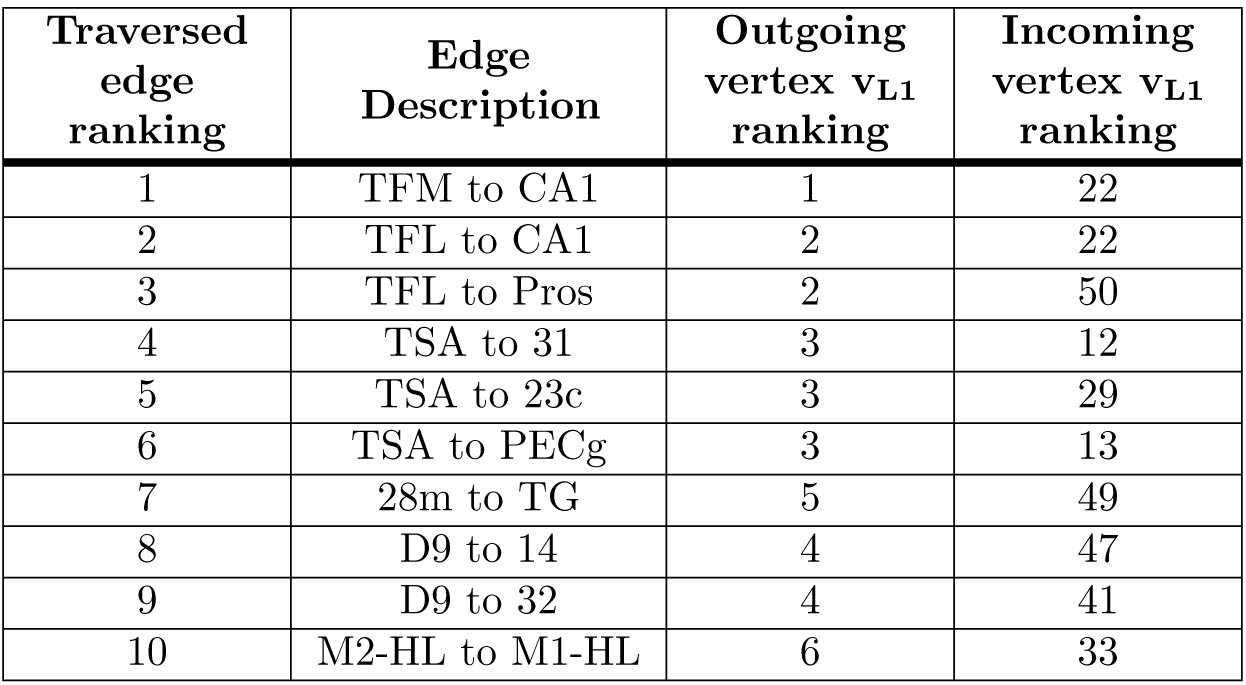
Most traversed edges with vertex *v*_*L*1_ rankings.

| Traversed edge ranking | Edge Description | Outgoing vertex $v_{L1}$ ranking | Incoming vertex $v_{L1}$ ranking |
| --- | --- | --- | --- |
| 1 | TFM to CA1 | 1 | 22 |
| 2 | TFL to CA1 | 2 | 22 |
| 3 | TFL to Pros | 2 | 50 |
| 4 | TSA to 31 | 3 | 12 |
| 5 | TSA to 23c | 3 | 29 |
| 6 | TSA to PECg | 3 | 13 |
| 7 | 28m to TG | 5 | 49 |
| 8 | D9 to 14 | 4 | 47 |
| 9 | D9 to 32 | 4 | 41 |
| 10 | M2-HL to M1-HL | 6 | 33 |

### Human Functional Connectome

The effectiveness of analysing brain connectomes, with network eigenvectors, has been demonstrated with the *C. elegans* and the macaque connectomes. The CDR shall now be employed on a series of connectomes generated by Roncal et al. [19] from magnetic resonance imaging (MRI) scans carried out by Landman et al. [2]. The voxels are defined as the intersection points on a three dimensional grid where each point is 1 mm apart from its neighbours. Each voxel is taken as a vertex in the network, resulting in a graph of 1,827,240 vertices. The edges of the network are undirected and defined as any two vertices that are connected by at least a single fibre where an edge weight of 1 represents a single fibre connection. This results in a network of weighted edges with some edges representing thousands of fibres connecting two regions.

Landman et al. scanned each subject twice, with a short break between scan and rescan [2]. Landman et al. used 21 healthy volunteers, but one of the voxelwise networks was unavailable for this paper therefore subject 127 is not included. The 20 remaining subjects are aged between 22 - 61 years with an even gender split. For each subject the first ten eigenvectors were analysed, those corresponding to the largest eigenvalues in magnitude of the adjacency matrix (equivalent to the smallest non-zero eigenvalues of the Laplacian matrix). A prominent pathway was detected for each of the ten first eigenvectors. The pathway was identified as the most prominent community according to the CDR approach. Prominence being determined by which community contained the vertex with the largest eigenvector entry in magnitude; referred to as the prominent vertex (PV).

### Subject 113

The eigenvector pathways of a 28 year old, right-handed, female (subject ID 113 [2]) are listed in Table 3. The top ten eigenvectors of scan 1 are displayed in the table alongside the closest matching pathways from scan 2. Fig. 4 displays only the matching pathway pairs if they achieved a percentage match of 60% or greater, therefore the pathways *v*_1_, *v*_2_, *v*_3_, *v*_4_, *v*_5_, *v*_7_ & *v*_10_ are displayed for scan 1 and *v*_2_, *v*_3_, *v*_6_, vg & *v*_10_ are shown for scan 2. Fig. 4 demonstrates that overlapping pathways may not include a similar number or distribution of vertices. This difference in length and density makes it more difficult to create an accurate metric for matching pathways. The metric developed here considers a threshold distance for all the vertices of one path to the nearest vertex in another and is described in more detail in the Materials and Methods Section. The percentage of matching vertices is detailed for subject 113 in Table 3. This table reveals that five of scan 2’s eigenvector pathways overlap with pathways in scan 1 where the majority of their vertices are neighbouring voxels. Table 3 also includes the distance between PVs from the pathways under comparison, which reveals that the majority of PVs are in close proximity as well.

**Table 3.** Comparison of first ten eigenvector pathways of scan 1 with the closest matching pathway from scan 2, for subject 113.

| Eigenvector |  | % match | PV Dist. [mm] |
| --- | --- | --- | --- |
| Scan 1 | Scan 2 |  |  |
| 1 | 8 | 76 | 2.2 |
| 2 | 2 | 100 | 9.4 |
| 3 | 8 | 94 | 2 |
| 4 | 10 | 61 | 3.7 |
| 5 | 3 | 87 | 3 |
| 6 | 10 | 22 | 8.7 |
| 7 | 8 | 76 | 3.2 |
| 8 | 1 | 0 | 43.3 |
| 9 | 1 | 0 | 13.3 |
| 10 | 6 | 60 | 3.3 |

**Fig 4.**
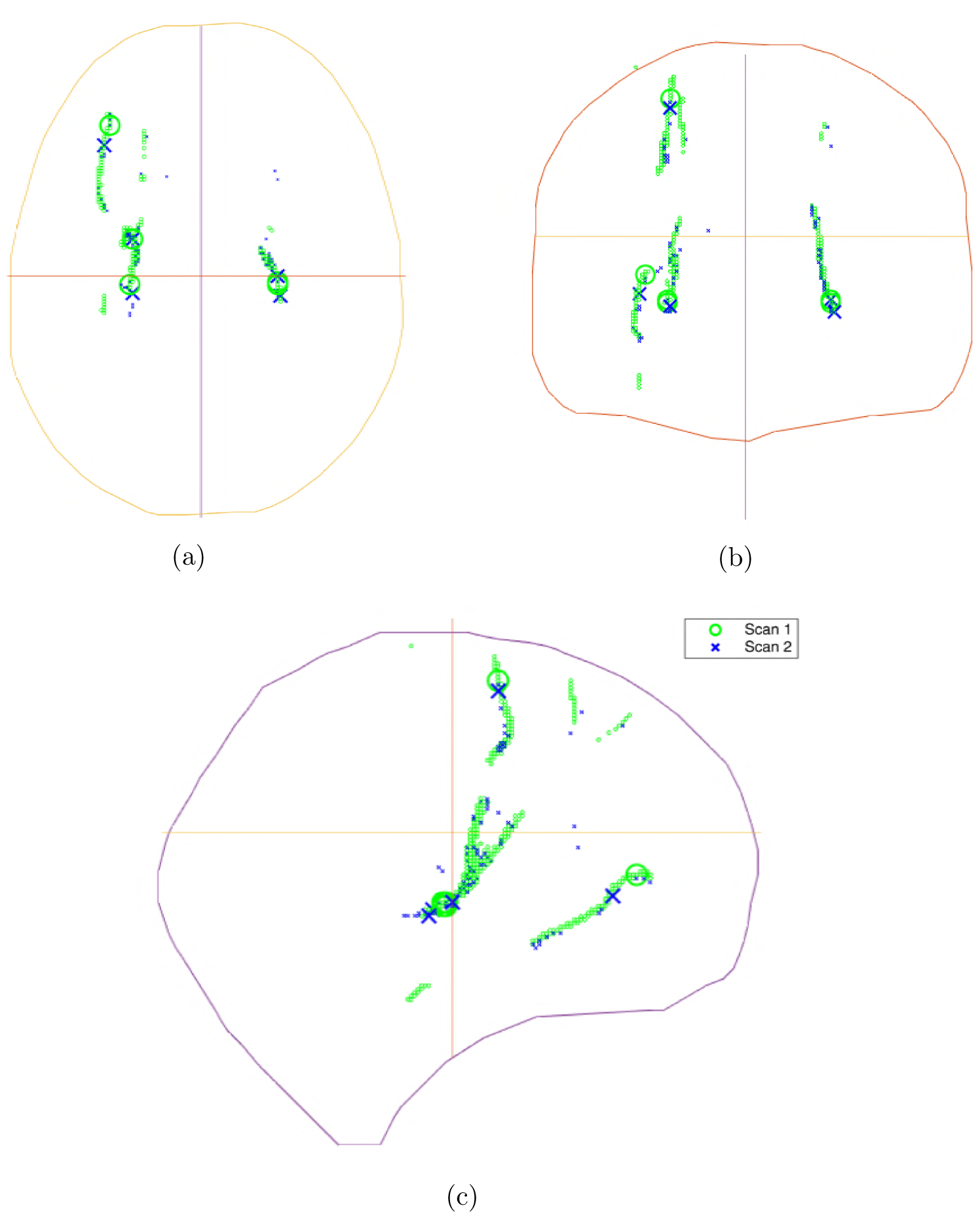
Human brain of subject 113 represented by x,y,z outlines from a brain surface model. Prominent matching pathways from scan 1 (*v*_1_, *v*_2_, *v*_3_, *v*_4_, *v*_5_, *v*_7_ & *v*_10_) and scan 2 (*v*_2_, *v*_3_, *v_6_ v*_8_ & *v*_10_) are displayed in green and blue respectively. The most prominent vertex for each eigenvector pathway is marked with a large circle or cross. (a) View from above; (b) View from behind; (c) View from the side with eigenvectors labelled.

The distribution of high weight edges in scan 1 and scan 2, for subject 113, are displayed in Fig. 5. The highest weighted edges are in a similar location for scan 1 and scan 2 with these locations also coinciding with the location of the top eigenvector pathways shown in Fig. 4. In spite of these similarities, a clear difference in the distribution and weightings of edges can be seen when comparing Fig. 5 (a) & (b).

**Fig 5.**
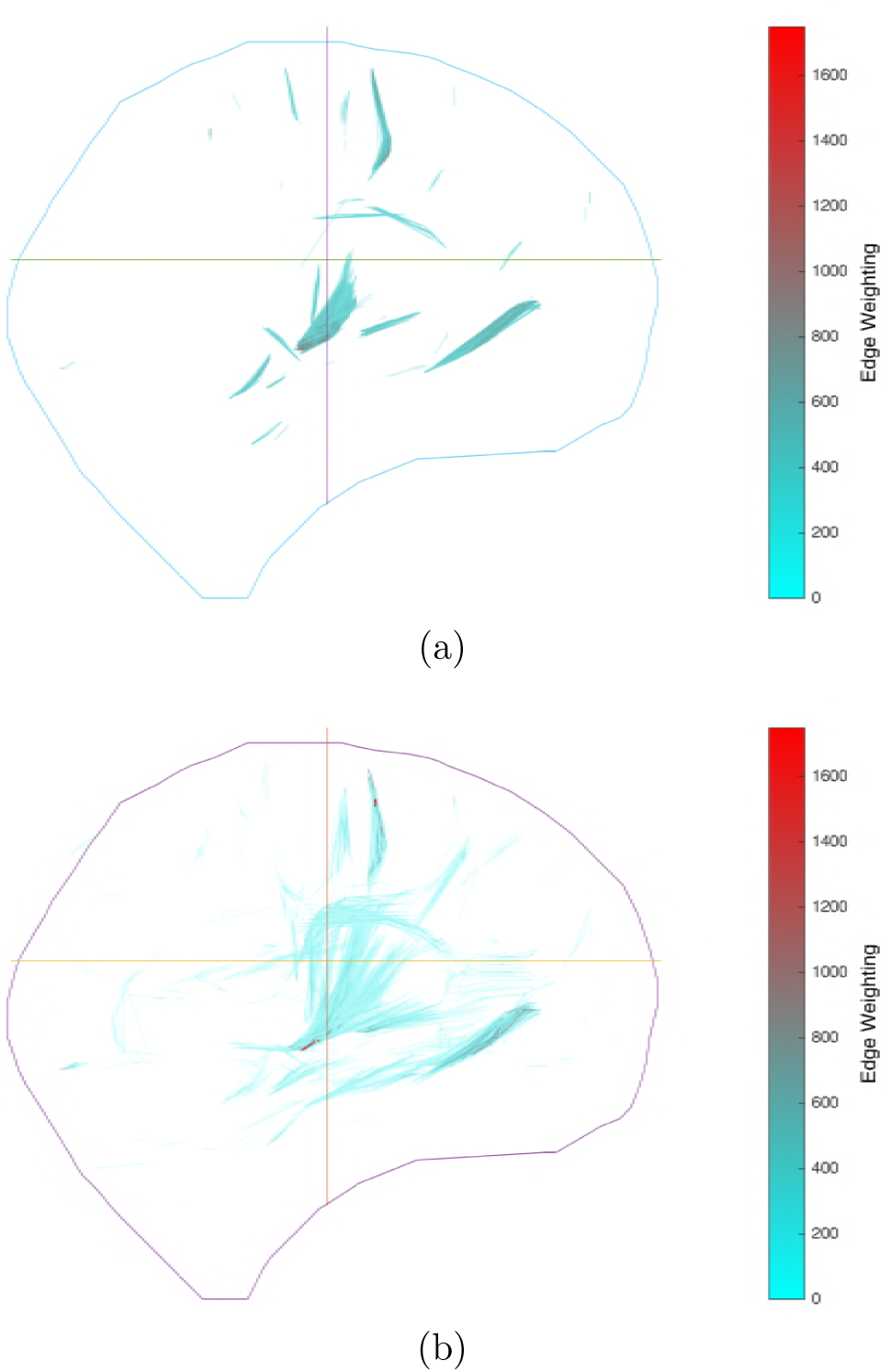
Brain of subject 113 represented by x,y,z outlines from a surface model. The 3500 highest traffic edges are displayed and coloured according to their weighting (a) View from side for scan 1; (b) View from side for scan 2.

These differences make subject identification difficult if considering only Fig. 5. However, the eigenvector pathways are not significantly affected by the change in edge weighting/distribution and, as a result, subject identification is possible with multiple pathways shown to be a high percentage match indicating that the scans belong to the same subject.

Whilst the differences highlighted in Fig. 5 did not affect shape and direction of the pathways, it did influence the number of vertices present in the pathways where there were less than a third the number of vertices in scan 2’s pathways than scan 1. This is a result of scan 2 producing few relatively high weighted edges (> 1400) and a majority of low weight edges (< 600), whereas scan 1 has a more even distribution with the highest weighted edge only ~ 1400 but with many edges above 1000. This uneven distribution translates into the eigenvector entries where scan 2 contains only a few high entry values whilst scan 1 has a more even distribution. The reduction in pathway size is, therefore, due to fewer vertices in scan 2 achieving the threshold eigenvector entry value to be included in the pathway, see the Materials and Methods Section.

### Subject Comparison

In the work by Roncal et al. [19] the Frobenius norm was used to demonstrate the similarity of the scan-rescan matrices. In Fig. 6 (a) the Frobenius norm of the difference between scan 1 and scan 2 graphs, referred to as the Frobenius Distance, is displayed for 20 subjects from Landman et al.’s study [2]. The most similar matrices are highlighted for each column where the majority of the matches are scan-rescan pairs for the same subject. When considering the most similar matrices for each row; the lowest Frobenius distance pair involved scan 2 of subject 113 for most of the scan 1 subjects, excluding subject 239, 422 & 742 that matched with their scan-rescan pair.

**Fig 6.**
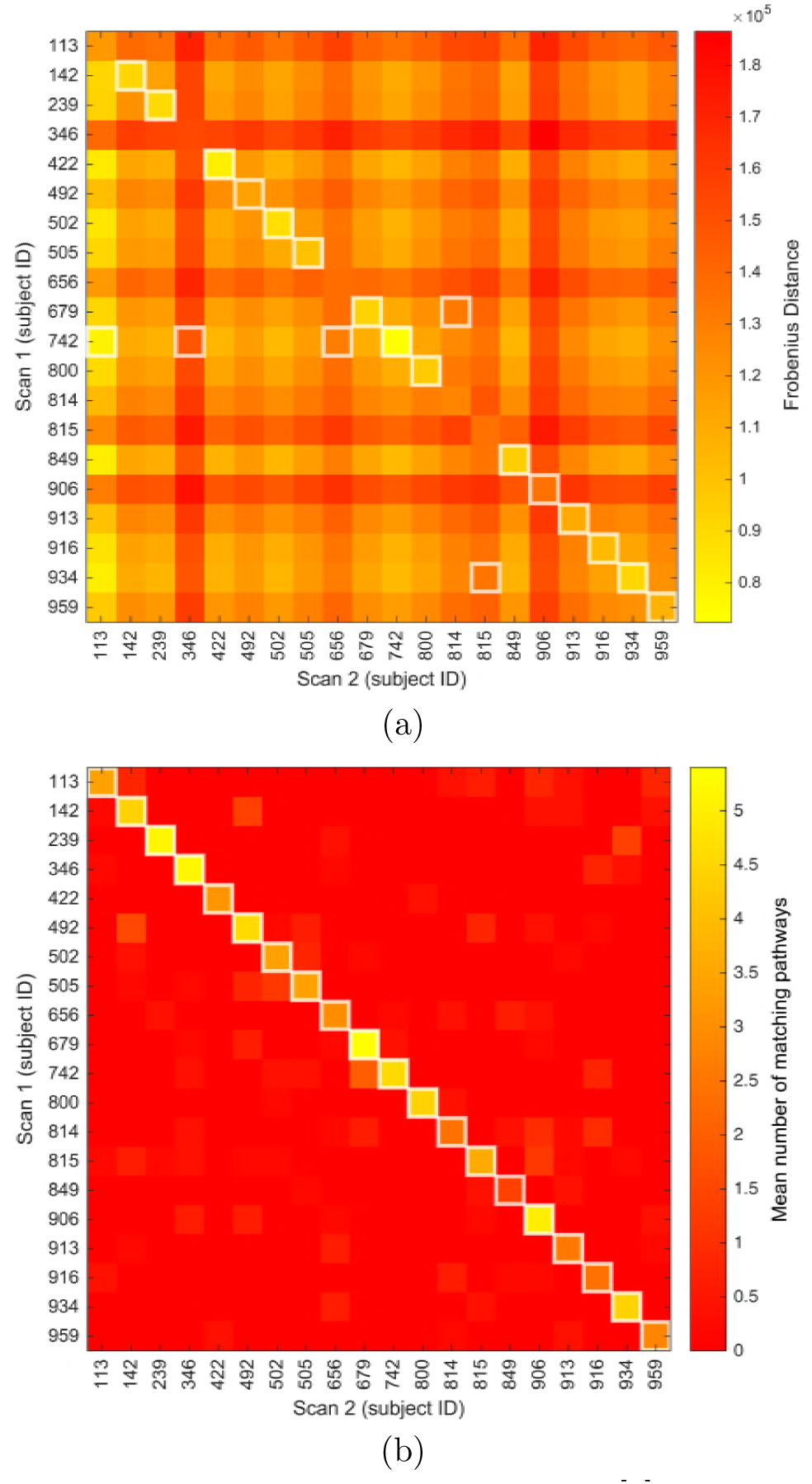
Comparison of scan 1 and 2 for Landman et al. [2] subjects where the most similar graph pairs are highlighted with a white outline. (a) Frobenius Distance where the most similar graph pairs are shown for each column (b) Mean number of matching pathways where the most similar graph pairs are for both row and column comparisons.

The eigenvector pathways were used in a similar manner to assess whether pathways matched, using the percentage match and PV distance categories shown in Table 3. Given the criteria in the Materials and Methods Section the mean number of matching pathways is presented in Fig. 6 (b). The Frobenius Distance can be seen in some cases to highlight a scan-rescan match, but the pathway matching approach is successful in every case with the matches clearly distinguished from non-matching pairs for both row and column comparisons.

The subject with the lowest number of matching pathways, as detailed in Fig. 6 (b), is subject 849. Subject 849’s best match is still the scan-rescan pair, but it also produced a lower mean number of matching pathways than some comparisons that were not scan-rescan pairs, such as the comparisons between subject 142 and subject 492. Investigating the pathways of subject 849 reveals that the shape and position of the pathways appear to be similar, see Fig. 7, which would have produced a high number of matching pathways. But the pathways are, with one exception, slightly offset from each other despite mostly matching in terms of shape, direction and length. It is this offset that reduces the mean number of matching pathways to 1.4 when visually there appears to be at least 4 matches, possibly 5. There are always errors in the images produced from MRI scans, even when using the same equipment and procedure, with small errors occurring because of slight changes in image orientation and magnetic field instability [20]. It is possible that these offsets are a result of such errors.

**Fig 7.**
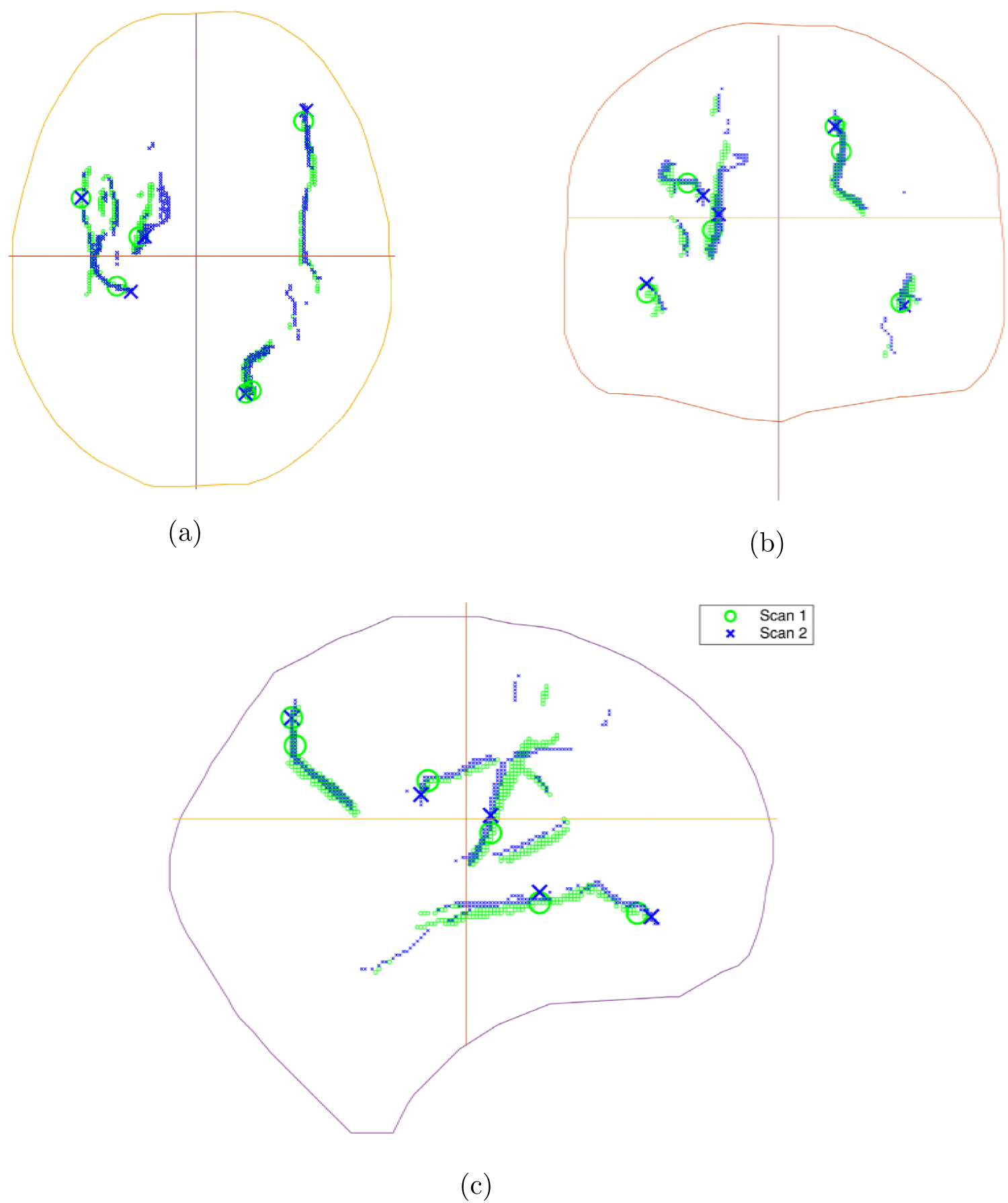
Human brain of subject 849 represented by x,y,z outlines from a brain surface model. Prominent vertices from scan 1 are displayed in green with scan 2 marked in blue. The most prominent vertex for each eigenvector is marked with a circle or cross. View from the side with eigenvectors labelled.

## Discussion

We found that incorporating multiple eigenvectors in the detection of communities produced a more nuanced picture of a system’s circuitry than had previously been achieved by using a single eigenvector. The communities detected are, due to their eigenvector-based nature, ordered by their effectiveness at channelling information to key vertices. For the small neuronal networks, of the *C. elegans* and the macaque, this problem is tractable for numerical flow simulations that model where and how information flows. Eigenvectors were able to produce similar findings to these numerical models where in the case of the macaque (CoCoMac graph) the hippocampus was demonstrated to be an influential region, despite previous claims that it was not possible to gain such information from analysing only the static network topology. This eigenvector approach was able to identify prominent pathways in large graphs, with millions of vertices, where numerical flow analysis is likely to be intractable. By comparing the number of matching pathways individuals can be identified from twenty subjects in the Landmann et al. scan-rescan dataset. This capability could have the potential to provide a quantative evaluation of how a subjects cognitive approach to a task changes over time. Since the most prominent pathways remain similar between scan and rescan, this method could assess if there were changes to the pathways used to accomplish the task every time the task was repeated.

The results of graph based analysis, and therefore the eigenvector method presented, is limited by the network that has been constructed. In particular, for human brains, undirected networks are the most prevalent but approaches do exist for the creation of directed connectomes from MRI scans. Without a directed network the insights available are limited as the true dynamics of the brain are concealed with key information lost by ignoring the imbalance in the outdegree to indegree ratio of vertices. It is known that the brain employs directed connections and without knowledge of these it is difficult to uncover if a prominent vertex is a sink or source of information in the graph. Indeed the Laplacian matrix, used for the analysis of the directed graphs, emphasises this imbalance further as each diagonal element is equal to the sum of the non-diagonal elements in its row i.e. the indegree of a vertex. But, as was displayed previously with the investigation of *C. elegans* and their electrical junction network (see the C. Elegans Connectome Section), insights can still be gained when using an undirected graph. As noted in the previous section, another source of uncertainty comes from the scan that generated the connectome where slight errors can affect the results, in particular the location of prominent pathways in the brain.

It is possible to conject as to how such connectome analysis could be used. Each pathway/community is associated with an eigenvector, the pathways are therefore ranked by their associated eigenvalue with the first eigenvector/eigenvalue associated with the largest dynamic response of the system. This provides a metric for ranking the prominence of the pathways in the brain. It is possible that such a capability could be applied as a quantitive assessment of, for example, a stroke victim’s progress in retraining neural pathways to regain speech. The prominence of the relevant neural pathways could be monitored to observe their growing prominence as the pathways are retrained. This could provide a metric with which to measure progress and provide further insight into the process of brain plasticity. This analysis could even form the basis of the treatment itself, where it has been observed that improved understanding of pathological circuitry has already guided deep brain stimulation used in the treatment of Parkinson’s disease, depression, and obsessive compulsive disorder [21]. There is also potential for employing noninvasive brain stimulation techniques, such as transcranial magnetic stimulation, as a significant part of the challenge is in identifying specific neural systems that should be targeted for intervention [21].

## Materials and Methods

A graph is defined as 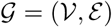, where there is a set of 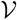 vertices and 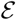 edges, which are unordered pairs of elements of 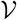 for an undirected graph and ordered pairs for a directed graph. The degree of a vertex is the number of edges connected to that vertex. In the case of a directed graph, there is an indegree and outdegree; indegree is the number of connections entering a vertex and outdegree is the number of connections exiting a vertex.

The adjacency matrix, *A*, is a square *n* × *n* matrix when representing a graph of *n* vertices. This matrix captures the network’s connections where *a_ij_* > 0 (*a_ij_* is the *ij*^th^ entry of the graph’s adjacency matrix) if there exists a directed edge from vertex *i* to *j* and 0 otherwise. Variable edge weights contain information on the relative strength of interactions, whilst uniform edge weighting either only represent the presence of a connection or is a result of all the edges having the same information carrying capacity. For an undirected graph, the adjacency matrix is symmetric with an edge 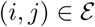 resulting in *a_ij_* = *a_ji_* > 0.

The Laplacian matrix is composed of the adjacency matrix and the degree matrix, *D*, as
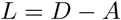

where the degree matrix is a diagonal matrix and the *i*^th^ diagonal element is equal to the outdegree of vertex *i*, which is equivalent to summing the elements of row *i* of *A*.

The eigenvectors of both the Laplacian and adjacency matrices are considered in this work. The dominant eigenvalue, *λ*_1_, for the adjacency matrix is the largest eigenvalue in magnitude while for the Laplacian matrix it is the smallest eigenvalue (*λ*_1_ = 0) [22]. The eigenvector associated with *λ*_1_ is referred to as the first eigenvector, specifically the first left eigenvector (FLE) when considering the Laplacian matrix of a directed graph.

The direction of an edge in this work defines the direction of travel for information. Therefore, if an edge is going from vertex *i* to vertex *j*, information is travelling from *i* to *j*. This is important as it affects the interpretation of the FLE. For example, if a packet of information departs every vertex in the network then the largest elements of the FLE are the vertices that information is funnelled towards. If the direction of information travel along an edge was reversed then the FLE would identify the vertices that are most effective sources for spreading information quickly across the whole network. This knowledge has been used previously to allocate resources that drive a network to a fast convergence to consensus [23], [24].

Considering the eigenvectors that proceed the first eigenvector, they can be understood to highlight vertices that collate information from across the whole network. But each proceeding eigenvector represents a slower mode of response for the system, therefore the vertices highlighted receive the information more slowly than the vertices that were prominent according to the first eigenvector or any associated with a smaller eigenvalue of the Laplacian matrix than the eigenvalue being considered.

### Communities of Dynamic Response

The Communities of Dynamic Response (CDR) algorithm detects communities that form in the presence of network stimulus. CDR is based on analysing three, usually consecutive, eigenvectors and is presented in detail in Algorithm 1. The algorithm works by assessing the coordinates for each vertex as defined by three chosen eigenvector entries. The most prominent vertices are located furthest from the origin of this eigenvector-based coordinate system and these vertices do not have an outgoing connection to a vertex that is at a greater distance from the origin. This can be seen in Fig. 8 (a) & (b) where communities are comprised of vertices that are each associated with one of the most prominent vertices (highlighted by a black outline). The communities are seen to spread from the origin of the plot out towards a prominent vertex.

**Fig 8.**
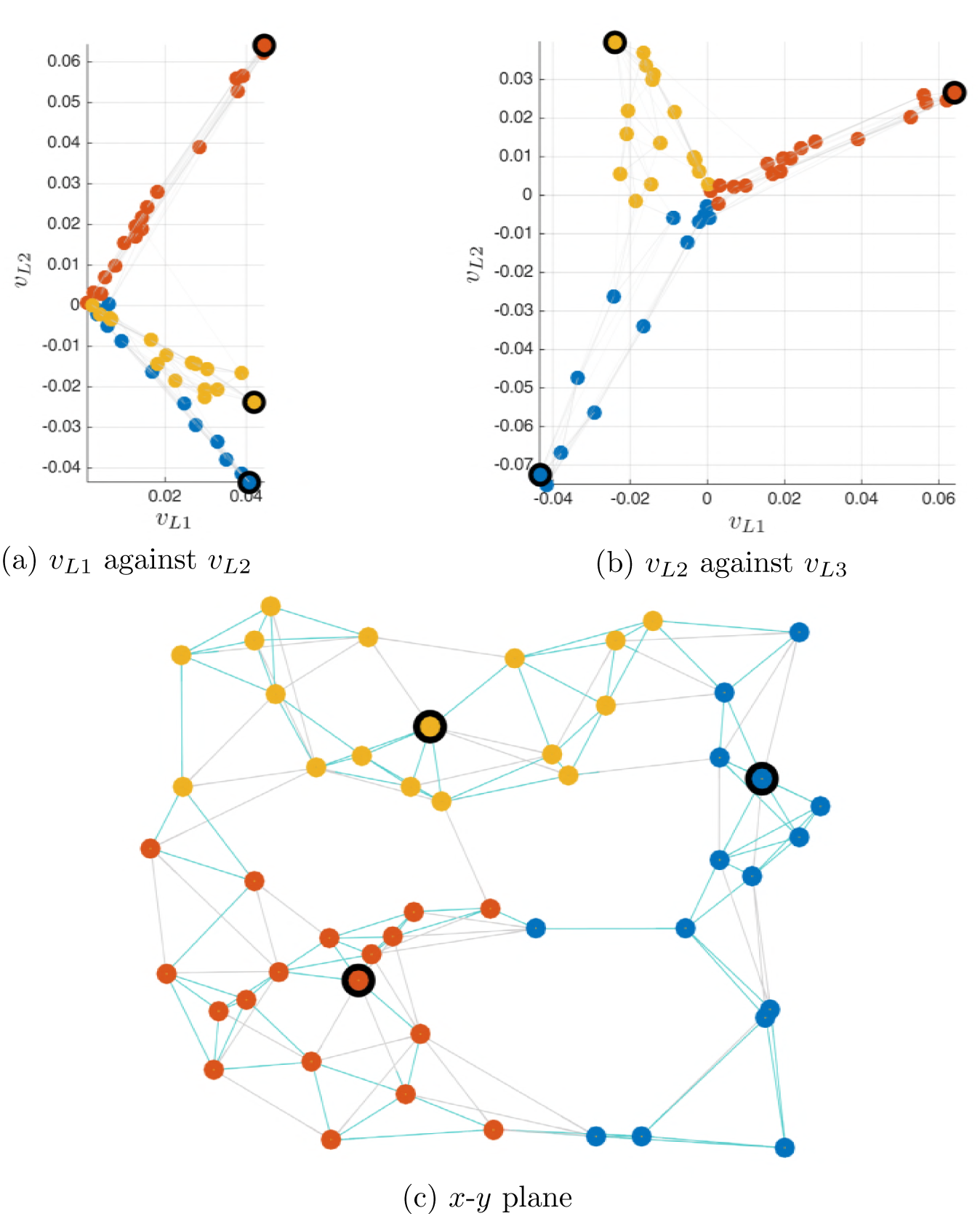
50 vertex, 5 outdegree, *k*-NNR graph with vertex colour indicating communities according to communities of dynamic response (Algorithm 1) and a black circle highlighting the most prominent vertex in each community. Visualisation according to (a) & (b) eigenvector space, where *v*_*L*1_, *v*_*L*2_ and *v*_*L*3_ are the first three left eigenvectors of the Laplacian matrix; (c) vertex position in *x-y* plane.

When considering the dynamics of the whole network, the FLE determines the most prominent vertices where, as was explained in the previous section, it represents the fastest response of the system. These prominent vertices in Fig. 8 are marked with a black outline and represent the nodes with the largest, in this example, *v*_*L*1_ value that are not connected to a node with a greater *v*_*L*1_ value.

The toy example in Fig 8 employs a *k*-Nearest Neighbour topology, whereby vertices are randomly distributed on a plane before connecting to their five nearest neighbours. The *k*-NNR topology is used since it is a good topology for demonstrating community structure as the nearest neighbour rule encourages communities to form.

#### Algorithm 1 Detecting communities of dynamic response

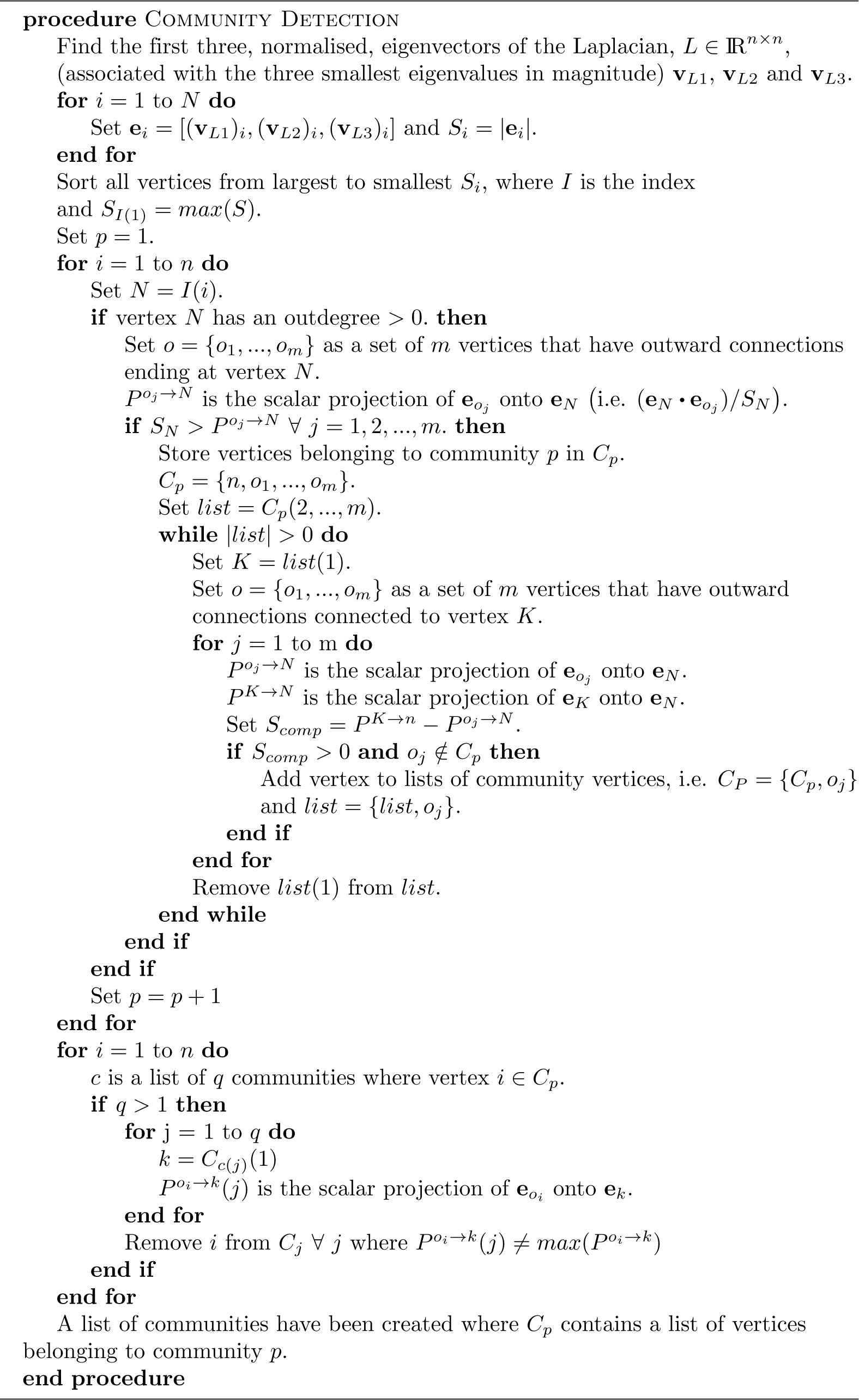

### Hippocampus in the Queueing Network

This section refers to the information flow model used by Mišić et al. [6] to investigate the CoCoMac connectome. The model was setup as a discrete-event queueing network where signals were continually generated, at randomly-selected grey matter vertices in the network, and assigned randomly-selected destination vertices. The signals then travelled through the network via white matter projections (edges). Grey matter vertices were modelled as servers with a finite buffer capacity, such that if a signal unit arrives at an occupied vertex, a queue will form. Upon reaching its destination vertex, the signal unit was removed from the network. Mišić et al. used this numerical model to show that the hippocampus (CA1) is a central hub. Demonstrating that the CA1 experiences a high throughput of signal traffic that places it in the top 3% for the total number of signal units that arrive at a vertex, the mean number of signal units at a vertex and the proportion of time a vertex is occupied by signals.

### Human Brain Pathways

#### Matching Pathways

For the analysis of human connectomes, a prominent pathway was detected for each of the ten first eigenvectors. Each pathway was selected as the most prominent community, from the communities generated by the CDR algorithm (see Algorithm 1). Prominence was determined based on which community contained the vertex with the largest eigenvector entry in magnitude; referred to as the prominent vertex (PV).

For a given eigenvector, *v*, only community nodes with min (*v*) > 0.01 were included in the pathway. This ensured that the pathways only included the most prominent members of each community, which produced clearer results when performing a pathway comparisons.

The metric developed considers the shortest distance from all the vertices of one path to the nearest vertex that belonged to the other path. Vertices were considered overlapping if they were from the same voxel or they were in an adjacent voxel (i.e. *<* 1.42 mm distance away). The percentage of vertices within this overlapping distance was then calculated. To determine if a pathway matched a threshold percentage match had to be achieved. For the results in this paper a mean value was taken for a range of threshold values. Therefore the mean number of overlapping pathways was checked for a range of percentage match thresholds from 50% to 90%, checked at 10% increments with a requirement that the PV distance be less than 15 mm. PV distance is a comparison of the point-to-point distance between the PV’s belonging to the matching pathways.

A pathway from one scan might be the closest match to multiple pathways from another scan, in this case only the closest match would count with the other matches ignored. For the example shown in Table 3, an eigenvector pathway has 7 matches with both 1, 3 and 7 from scan 1, therefore two of these would be ignored for a 50% matching criteria resulting in 5 matching pathways in total.

#### Frobenius Distance

When applying the Frobenius norm to the difference between two matrices it is often referred to as the Frobenius distance and defined as
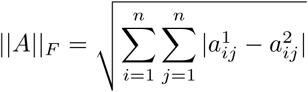

where *a*^1^ and *a*^2^ are elements of the adjacency matrix for scan 1 and scan 2 respectively.

